# Discovering genomic islands in unannotated bacterial genomes using sequence embedding

**DOI:** 10.1101/2022.08.25.505341

**Authors:** Priyanka Banerjee, Oliver Eulenstein, Iddo Friedberg

## Abstract

**Motivation:** Genomic islands (GEIs) are clusters of genes in bacterial genomes that are typically acquired by horizontal gene transfer. Genomic islands play a crucial role in the evolution of bacteria by helping them adapt quickly to changing environments. Specifically of interest to human health, many GEIs contain pathogenicity and antimicrobial resistance genes. Detecting GEIs is therefore an important problem in biomedical and environmental research. There have been many previous studies for computationally identifying GEIs, but most of the studies rely either on detecting differences between closely related genomes, or on annotated nucleotide sequences with predictions based on a fixed set of known features.

**Results:** Here we present TreasureIsland, which uses a new unsupervised representation of DNA sequences to predict GEIs. We developed a high precision boundary detection method featuring an incremental fine-tuning of GEI borders, and we evaluated the accuracy of this framework using a new comprehensive reference dataset, Benbow. We show that TreasureIsland performs competitively when compared with other GEI predictors, enabling the identification of genomic islands in unannotated and taxonomically isolated bacterial genomes.

**Availability:** The source code and the datasets used in this study are available at: https://github.com/priyamayur/GenomicIslandPrediction

**Supplementary information:** Supplementary Material is available at *Bioinformatics* online.

## 1 Introduction

Horizontal gene transfer (HGT) in bacteria is a major mechanism for acquiring genetic material, enabling adaptation to a changing environment by rapidly conferring new phenotypes, such as stress resistance and antibiotic resistance [1, 2]. Genomic islands (GEIs) are clusters of genes acquired by HGT, providing evolutionary diversity and by conferring complex traits requiring several gene products that are co-expressed are rapidly put into play [3–7]. GEIs are typically classified based on their functional content: pathogenicity islands that contain pathogenic or virulent genes, resistance islands containing antimicrobial-resistant genes, symbiosis islands containing genes that establish symbiosis with host organisms, or metabolic islands containing adaptive metabolic capabilities. (For review see: [8]). GEIs have some distinguishing features, including (i) a typical size range of 10-200 kbp [9], (ii) a sequence composition that is generally different than the core genome, and (iii) frequent associations with tRNA-encoding genes, flanking direct repeats and mobility genes, with a high prevalence of phage related genes and hypothetical proteins [5]. The wide range of adaptive functions makes the identification of GEIs of particular environmental and biomedical interest [3, 9].

GEIs are identified experimentally using methods such as DNA-DNA hybridization, subtractive hybridization, or using counter-selectable markers [5, 10, 11]. However, experimental methods are limited to certain combinations of bacterial strains and GEI types, and can be expensive and time-consuming. Therefore, reliable computational GEI prediction methods are needed.

Methods for computationally predicting GEIs are broadly divided into two approaches: comparative genomics, and sequence composition. Comparative genomics approaches involve the use of closely related bacterial and archaeal genomes [8]. A GEI is identified when a cluster of genes is present in an organism that is not present in any related genomes [12]. Recently, Bertelli and colleagues conducted research showing that comparative genomics based approaches can predict GEI boundaries accurately. However, such methods depend on the availability of closely related genomes, and results vary widely based on which genomes are selected [8]. Sequence-composition methods are based on identifying atypical subsequences in the chromosome. These methods identify aberrations in structural features, such as GC content, dinucleotide content, codon usage, k-mer count, presence of different insertion sites, the presence of mobility genes, phage genes, hypothetical proteins, and direct flanking repeats [3, 5]. Prediction methods that only use sequence composition are usually less accurate than those using annotated sequences and comparative genomics. This is due to the smaller feature set the sequence composition methods are working with [8]. In sum, there are certain limitations to current GEI prediction methods: (i) the requirement of closely related genomes in case of comparative genomics-based approach; (ii) dependency on annotated genomes and, (iii) lack of a good feature set in a sequence composition based approach.

Here we present TreasureIsland, a genomic island prediction software that uses an unsupervised representation of DNA sequences, and therefore does not require any computation of a fixed number of features. TreasureIsland does not require annotated genomes, nor does it require the availability of closely related genomes for reference. Specifically, TreasureIsland uses word embedding for the detection of differential sequence compositions. Word embedding methods such as Word2vec are particularly powerful for natural language processing as they capture the semantic meaning and the context of the words [13]. Word embedding has been used in several other bioinformatics applications, including novel ORF identification [14], DNA origin of replication [15], assignment of function to protein domains [16], mapping the gut microbiome [17], and protein family classification [18]. To the best of our knowledge, this is the first time word embedding has been used to discover genomic islands. In the era of high throughput sequencing of microbial genomes and metagenome assembled genomes (MAGs), such a tool is a needed addition to the arsenal of tools used for genomic island discovery.

## 2 Methods

Figures 1 & 2 provide an overview of TreasureIsland. There are two phases: first, the model construction, or GEI classification phases, involving embedding of DNA sequences. The second phase involves the identification of initial GEI segments using the model constructed in the first phase, followed by a refinement algorithm Figure 4 to identify the exact GEI borders.

**Figure 1:**
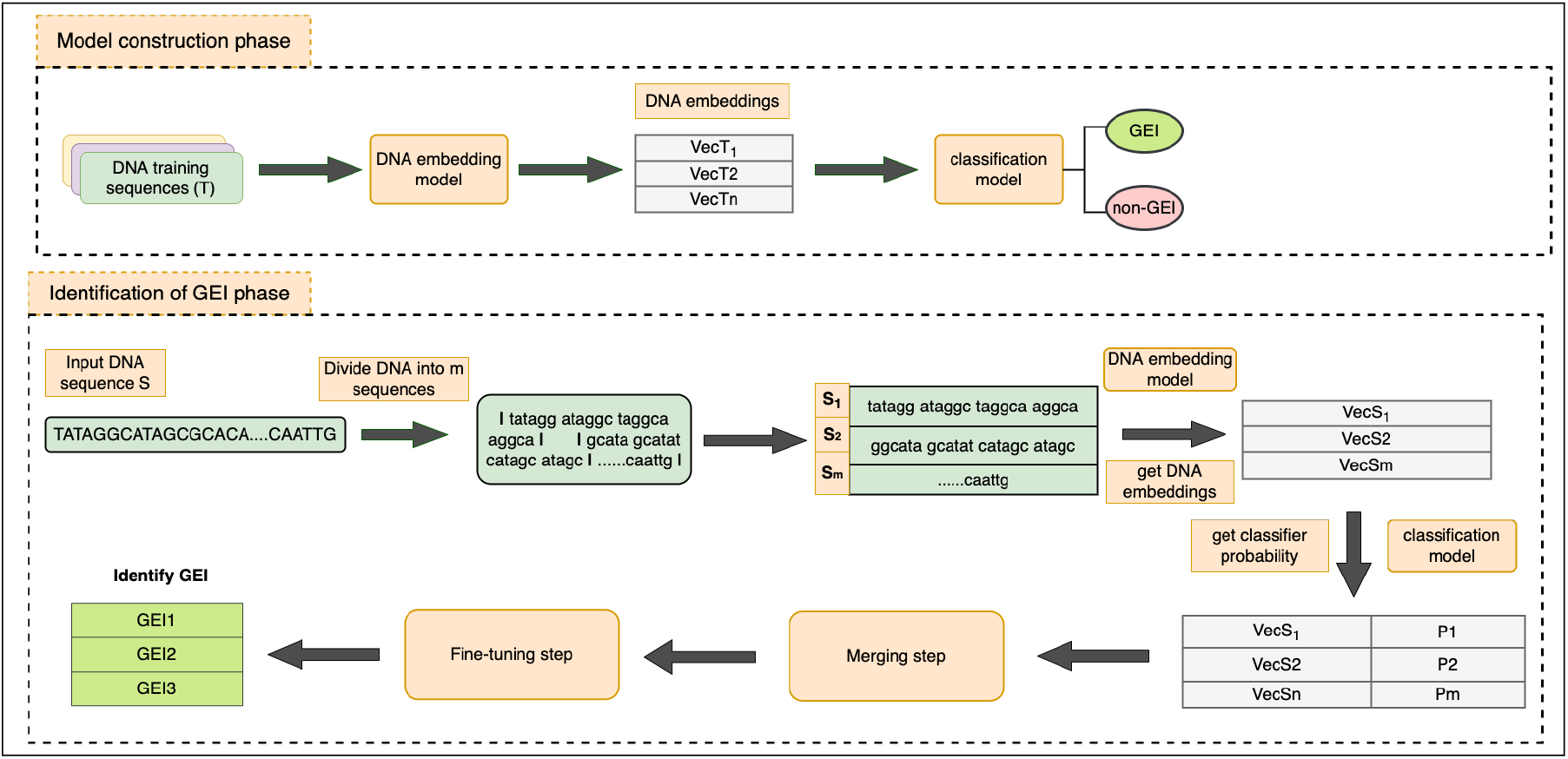
An overview of Treasurelsland. The Treasurelsland workflow consists of two phases: (i) The model construction phase for classifying DNA segments to GEI or non-GEI; this includes Doc2vec embedding of DNA sequences, and training a classifier. (ii) Identification of GEI location in the bacterial chromosome, which includes a fine-tuning of the GEI borders.

**Figure 2:**
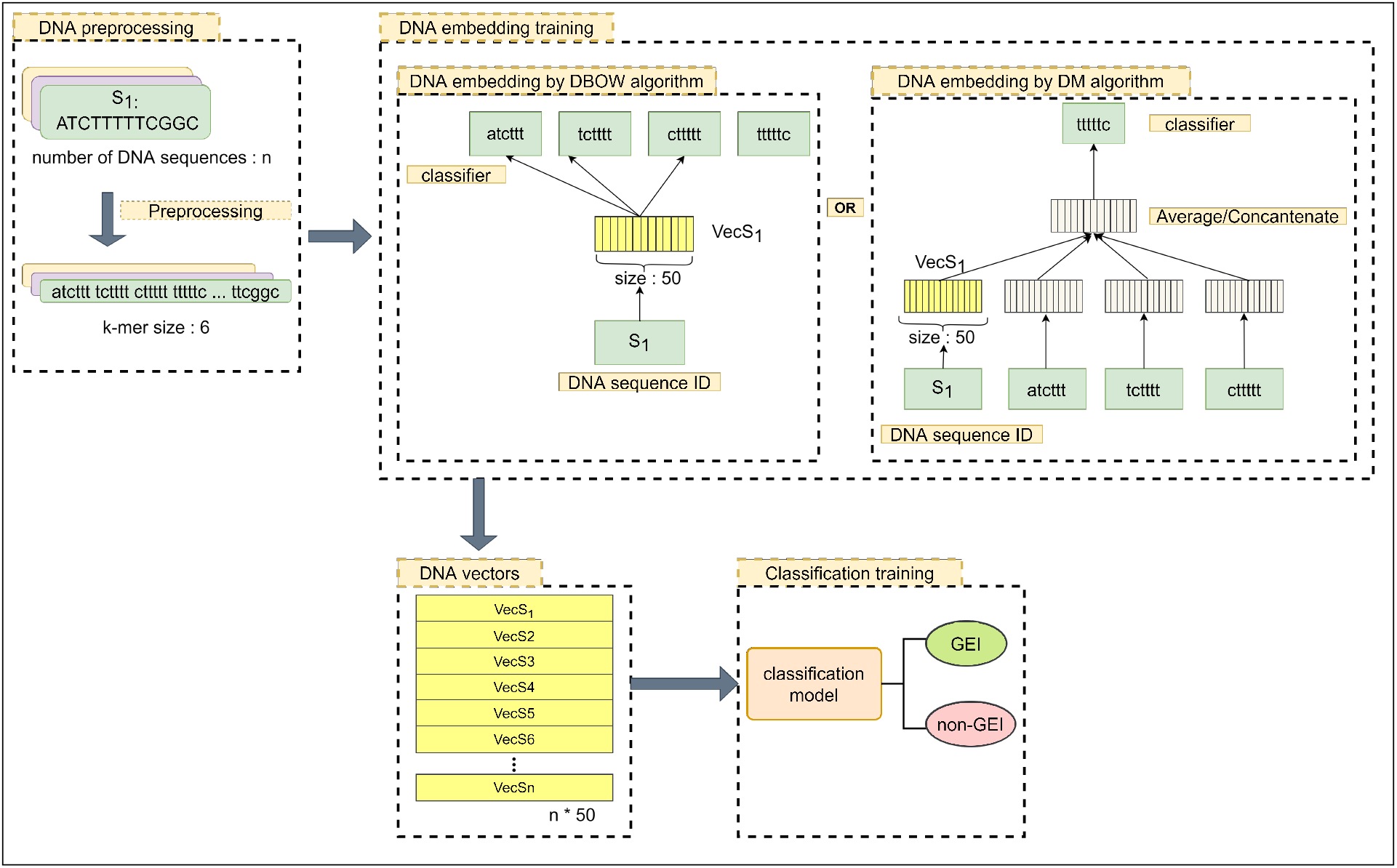
Model construction phase. Each DNA sequence in the training or validation set was first preprocessed and converted into a fixed length vector using either the Distributed Bag of Words (DBOW) or the Distributed Memory (DM) algorithm. These DNA vectors were then classified into GEI or non-GEI.

**Figure 3:**
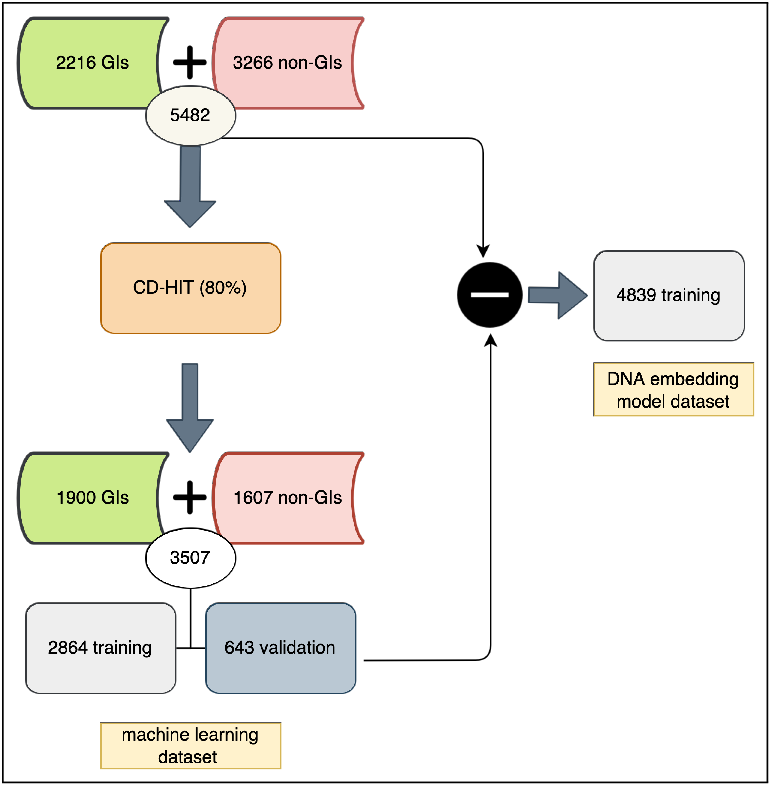
Creating the dataset for the model construction phase. The initial set of GEIs and non-GEIs was run through CD-HIT at 80% similarity to reduce redundancy and bias in the dataset. The machine learning validation data was formed by separating 20 genomes with 643 examples from a total of 212 genomes. To reduce bias in the DNA embedding model dataset, it was formed by removing the machine learning validation data from the initial GEIs and non-GEIs.

**Figure 4:**
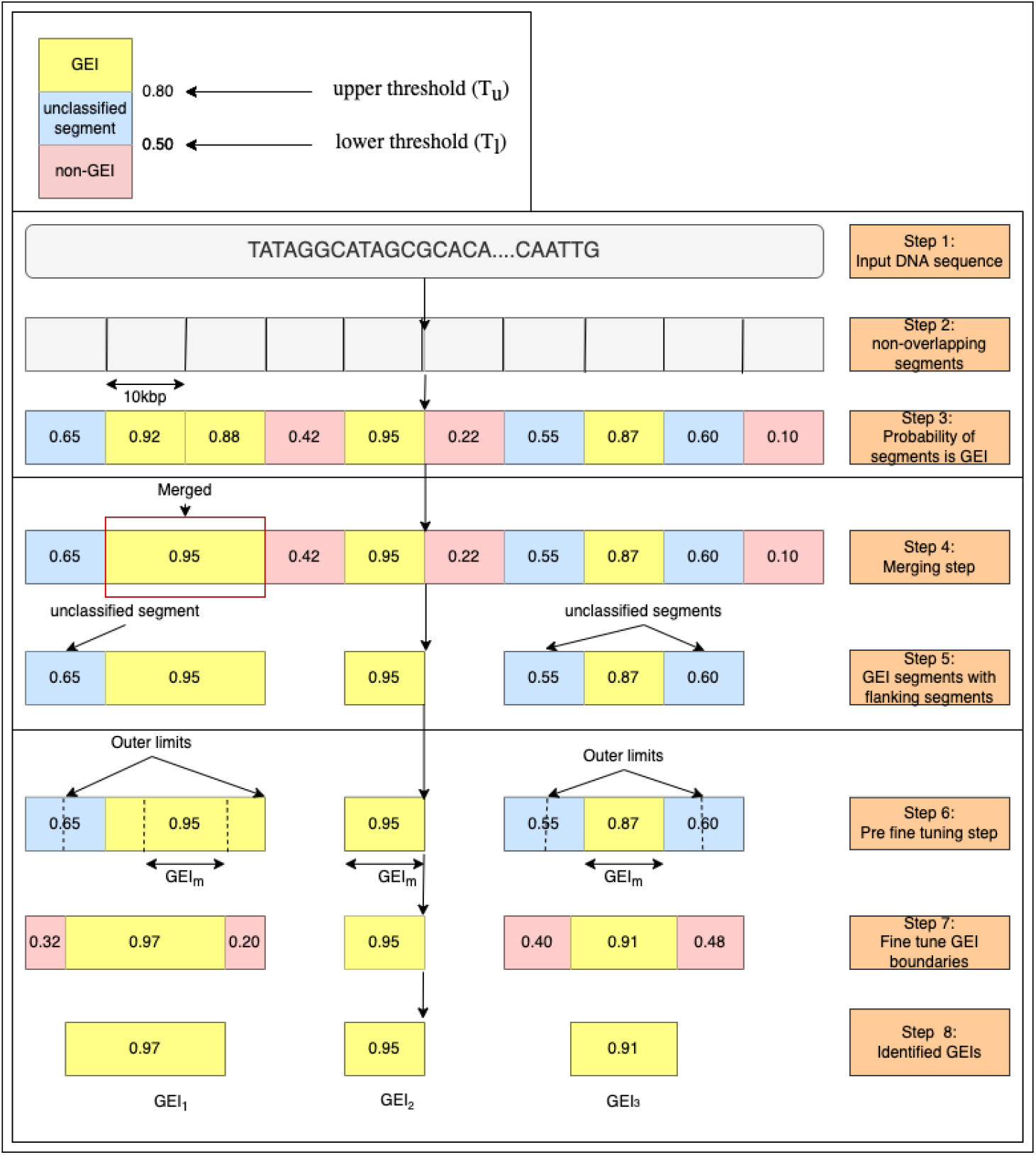
Identification of GEI phase. In this example, *T_u_* is set to 0.80, and *T_l_* is set to 0.50. The input DNA sequence is divided into non-overlapping DNA segments. The probabilities of each of the segments being a GEI are determined. Next, the adjacent positive GEIs are merged, and the unclassified segments are attached. These regions are then fine-tuned to find the final GEIs.

### 2.1 Dataset Construction

#### The Benbow dataset

To train and validate our model, we needed a large, accurate, and non-redundant dataset of genomic islands. The dataset should contain the islands within a whole genome context, and have a large non-redundant set of positive and negative examples. To create a large GEI data set for training TreasureIsland, we compiled data from four well-established GEI databases into a unified, non-redundant dataset we named Benbow. The datasets we used are listed in Table 1. The Main database (M) [19] incorporates the L database [12] that includes negative examples from literature curation. We also added the non-redundant (based on sequence similarity) from PAIdb (P) [21] to create the full database of positive and negative examples as elaborated. Benbow consists of GEI sequences (positive labels), which we call *BENBOW_pos_*, and non-GEI sequences (negative labels) which we call *BENBOW_neg_*. *All of BENBOW*: *BENBOW_pos_* ≔ *M_pos_* + *E* − (*E* ⋂ *M*) + *P* - (*P* ⋂ *M*) *BENBOW_neg_* ≔ *M_neg_*

**Table 1:** Sources of GEIs used to construct Benbow, the unified GEI set used in this study. See text for details.

| Notation | Total genomes | Number of GEI | Number of non-GEI | Data Source |
| --- | --- | --- | --- | --- |
| M (Main) | 104 | 1845 | 3266 | [19] |
| E (Early) | 32 | 269 | 0 | [20] |
| L (Literature) | 6 | 80 | 0 | [12] |
| P (PAI) | 105 | 264 | 0 | [21] |
| Benbow (M+E+P) | 219 | 2,216 | 3266 | - |

To construct *BENBOW_pos_* we added 1. GEIs from all genomes in *M* (104 genomes), 2. GEIs from all genomes in *E* that do not overlap with the genomes in *M* (32 − 8 = 24 genomes, containing 172 GEI sequences) 3. GEIs from all genomes in *E* that do not overlap with the genomes in *M* (105 − 14 = 91 genomes, containing 199 GEI sequences). Thus, the total *BENBOW_pos_* data set is combined from 1, 845 (*M* dataset) + 172 (*E* dataset) + 199 (*P* dataset) = 2, 216 GEIs. The total *BENBOW_neg_* dataset contains 3,266 non-GEIs from all genomes in M. To remove redundancy and reduce bias, we ran CD-HIT [22] using an 80% sequence identity cut-off on the positive and negative label datasets. This resulted in the positive and negative examples of 1,900 sequences from 210 genomes and 1,607 from 99 sequences, respectively, as shown in Figure 3.

##### BENBOW machine learning dataset

Since the goal of our model is to predict GEIs within the context of a bacterial genome sequence, we created the training and validation datasets from whole genomes, rather than use “disembodied” positive and negative GEI & non-GEI sequences. To create the validation dataset, we randomly selected 20 genomes, with 399 GEIs and 244 non-GEIs (a total of 643), and for the training and tuning set we used 1,501 GEIs from 190 genomes and 1,363 non-GEIs from 79 genomes for a total of 2,864 sequences. To tune the parameters in the machine learning models, we used 10×cross-validation.

##### BENBOW DNA embedding model dataset

The unsupervised DNA embedding model needs a large redundant data set to understand the underlying patterns in the DNA sequences. Therefore, for the DNA embedding model dataset we used the dataset obtained before CD-HIT was applied and removed the machine learning validation set from it. This gave us a positive dataset of 1,817 GEIs and a negative dataset of 3,022 non-GEIs.

### 2.2 Computational Framework

The TreasureIsland computational framework consists of two phases: (i) the model construction phase for classifying DNA segments as GEI / non-GEI; (ii) GEI identification for any input sequence, typically a whole bacterial chromosome. As seen in the overview in Figure 1, in the model construction phase we construct an embedding model, which represents the variable-length DNA represented by fixed-length vectors. We then use these vectors to classify the segments of DNA into a GEI or a non-GEI region in a genome. At the end of the first phase, we are left with an embedding model and a classifier for DNA segments.

In the second phase, identification of GEIs, we divide an input sequence into nonoverlapping segments (Table 2, `sequence_window_size`). These segments are then embedded and classified using the embedding and classifier models, respectively, from the previous phase. The GEI classified segments are then processed to refine the boundaries to output the GEI regions in the input DNA sequence.

**Table 2:**
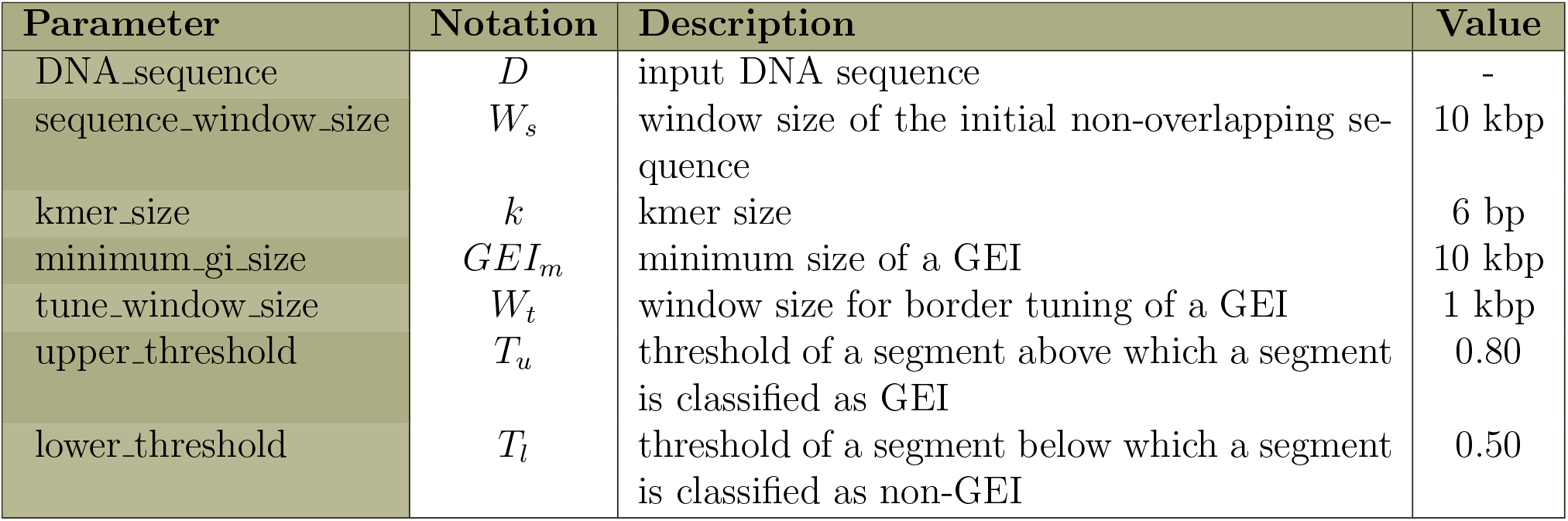
Parameters used for the GEI identification phase. See text for details.

| Parameter | Notation | Description | Value |
| --- | --- | --- | --- |
| DNA_sequence | $D$ | input DNA sequence | - |
| sequence_window_size | $W_s$ | window size of the initial non-overlapping sequence | 10 kbp |
| kmer_size | $k$ | kmer size | 6 bp |
| minimum_gi_size | $GEI_m$ | minimum size of a GEI | 10 kbp |
| tune_window_size | $W_t$ | window size for border tuning of a GEI | 1 kbp |
| upper_threshold | $T_u$ | threshold of a segment above which a segment is classified as GEI | 0.80 |
| lower_threshold | $T_l$ | threshold of a segment below which a segment is classified as non-GEI | 0.50 |

#### 2.2.1 Model construction phase

Here we construct the embedding and classifier models for DNA sequences.

##### DNA as a document

As noted in the Introduction, word embedding has been successfully adopted in bioinformatics to classify biological sequences in many applications. Doc2vec is an extension of the Word2vec model [23]. Doc2vec converts a variable-length document into a fixed-length vector; each document is then identified by a document ID and is then converted to a vector to represent every document. There are two different types of Doc2vec embedding models-Distributed Memory (DM) and Distributed Bag of Words (DBOW). In the DM model, a document ID is added as another word in addition to the words in the document. The DM model learns the word vectors along with the document vectors, which is done by predicting any given word using both the context words and the document ID. This model is analogous to the continuous bag of words (CBOW) model in Word2vec. In contrast, DBOW ignores the order of the words, and it predicts a randomly sampled word from the document given the document ID. This process is analogous to the Skipgram model in Word2vec.

##### Preprocessing DNA

For our purpose, any given sequence (GEI or not GEI) is considered a document and is represented as a sequence of overlapping *k*-mers which are the words. As shown in Figure 2, each GEI and non-GEI example (range of 1,300-65,000 bp each), which contains *k*-mers as its words, is tagged by a unique DNA sequence ID. We then trained two different document vector models: a Distributed Memory (DM) model, and a Distributed Bag of Word (DBOW) model.

##### Constructing the classifier

After we trained the embedding model, we obtained the vectors for the training set and validation set by using gradient descent from the embedding model, while fixing the rest of the model parameters as in [23]. We then fed the training vectors into several different supervised machine learning algorithms, to complete a binary classification task as GEI (Class 1) or non-GEI (Class 0).

##### Tuning hyperparameters for the model construction phase

The hyperparameters include the length of *k* and the choice between overlapping and non-overlapping *k*-mers. We tested all *k* ∈{3, …, 9}, and the optimal value was 6 with overlapping *k*-mers. At the document embedding level, the tuned hyperparameters are vector size, window, epoch, alpha values, dbow_words for document vector DBOW model, and dm_concat for document vector DM model. The best choice for the model was the DBOW model, and the optimal hyperparameters were found to be vector size 50, window 10, epochs 150, alpha 0.025, dbow_words 0. The classification task was selected to be performed by SVM, whose hyperparameters were found to be optimal at C=2, gamma = 1, kernel= RBF. These we obtained after tuning the model using a 10× cross-validated grid search results.

#### 2.2.2 Identification of GEI phase

This phase takes as an input a DNA sequence, usually a whole chromosome, and identifies all possible genomic islands in the sequence. The DNA-embedding model and the classifier from the first phase are used here. The parameters used for this phase are explained in Table 2.

##### DNA vectors representation

We divide the full chromosome sequence *D*, is divided into non-overlapping segments of fixed length. *D* = [*d*_1_, *d*_2_, *d*_3_, …, *d_n_*]. These segments are analogous to a document. We then take each of these segments (“documents”) and preprocess in the same way as the model construction phase by finding all *k*-mers of size *k*. The segments are then embedded by inferring vectors from the DNA-embedding model. Finally, we feed the vectors into the classifier and determine the probability (or confidence) *p*_1_, *p*_2_, *p*_3_, …, *p_n_* with *p_j_* = (0, 1], *p_j_* ∈ (0, 1] where 1 ≤ *j* ≤ *n* for class 1 (GEI) for each of the segments.

##### Merging

In the merging step, we merge adjacent segments identified as GEIs, where appropriate (Figure 4). We set two GEI probability thresholds to determine merging: an Upper Threshold *T_u_* and a Lower Threshold *T_l_*). Any segment with a predicted GEI probability *P*(*GEI*) having *P*(*GEI*) ≥ *T_u_* is labeled as a GEI. If two or more adjacent segments are found to be greater than or equal to *T_u_*, the segments are merged, and the merged sequence is considered to be a GEI (see Figure 4, Step 4, red box)/ Segments with *P*(*GEI*) ≤ *T_l_* are labeled as non-GEI. We considered the segments having probabilities between *T_u_* and *T_l_* unclassified, since we are unsure to which class they belong. To identify a more precise border, the unclassified segments are subject to the following procedure to eliminate them, and refine the GEI/non-GEI classification.

Each GEI segment uses Algorithm 1 to tune its left and right border. Step six shows the constraints used to tune the segments. The outer limit is set at the middle of the unclassified segment. This is to prevent an overlap of the GEI regions if there is another GEI on the other side of the unclassified segment. If a GEI has no unclassified segment, the outer limit is the original GEI border. The GEI segments cannot be lesser than the minimum GEI size considered *GEI_m_*.

Step seven shows how we tune borders on both ends of GEI, namely left and right borders. Algorithm 1 takes as input each border along with the presence of the unclassified segment on these borders and computes the optimized border position. It first determines the direction in which this border should shift, depending on the presence of the unclassified sequence. For instance, if we take as an input a left border with an unclassified segment, the fine-tuning direction is set to shift to the left. Next, the border is shifted in that direction by the tuning window size *W_t_* set in the parameter table, and on each shift, the probability for the current fragment to be a GEI is noted. This process continues iteratively and is stopped when the previous probability is greater than the current probability, or when any of the constraints are not satisfied. The optimized position for the border set as input is then obtained from the output tuple. The process enables us to reach a good balance between sensitivity and specificity for predicting GEI segments.

#### Algorithm 1: GEI border fine-tuning algorithm

This algorithm processes each border(left or right) of a GEI segment at a time

**Input:**

- *border*(*variable*) = (1, 0) denotes a *left* border and *border*(*variable*) = (0, 1) denotes a *right* border
- *has_unclassified_segment*(*variable*): boolean denoting the presence of an unclassified segment on either side of the known GEI
- *start, end*(*variables*): current start and end position of the GEI (Figure 4, yellow box)
- *outer_limit_start_, outer_limit_end_*(*variables*): start and end limits of the maximal extension or contraction of the GEI
- *d_left_, d_right_*(*variables*): direction in which the left and right borders should shift, respectively
- *GEI_m_*(*variable*) = 10kbp (from Table 2)
- *tune_window size* = *W_t_*(*variable*) = 1*kbp* (from Table 2)
- *FindGEIProbability*(*function*): function that finds the probability of a region to be GEI, given a start and end position.
- *FineTuneSet*(*object*): object representing ‘Start’, ‘End’, ‘*Probability_GEI_*’ for a segment.

**Output:** A *FineTuneSet* result with optimized border position (in bp) and probability of the segment is GEI `/* value of −1 sets the direction of the border to the left, and the value of 1 sets the direction of the border to the right` */

**if** *has_unclassified_segment* **then**

*d_left_* = border [0] × −1
*d_right_* = border [1] × 1

**else**

*d_left_* = border [0] × 1
*d_right_* = border [1] × −1

**end**

/* `for the left border`, *d_right_* `becomes fixed to 0 and for the right border`, *d_left_* `becomes fixed to 0` */

*probability ← FindGEIProbability*(*start, end*)

*current* ← *next* ← *FineTuneSet*(*start, end, probability*)

**while** *(next.Probability_GEI_ ≥ *current.Probability_GEI_* ***and*** *next.Start* ≥ *outer_limit_start_* **and***

*next.End* ≤ *outer_limit_end_* ***and*** *next.End* – *next.Start* ≥ *GEI_m_)* ***do***

*current* ← *next*
*next.Start* ← *next.Start* + *W_t_* × *d_left_*
*next.End* ← *next.End* + *W_t_* × *d_right_*
*next.Probability_GEI_* ← *FindGEIProbability*(*next.Start,next.End*)

**end**

**return** *current*

##### Hyperparameters for the GEI identification phase

The parameters required to identify the GEIs in the fine-tuning step are described in Table 2. Those include: windowsize, kmer_size, minimum_gi_size, tune_window, upper_threshold (*T_u_*) and lower_threshold (*T_l_*). These hyperparameters were obtained after tuning the identification of GEI results on 20 genomes by grid search. The grid search results were found to be optimal with window_size 10,000, kmer_size 6, minimum_gi_size 10,000 in keeping with the previous research on genomic island sizes [9], tune_window 1,000, upper_threshold (*T_u_*) 0.80, lower_threshold 0.5.

### 2.3 Evaluation of Treasurelsland’s Performance

To evaluate TreasureIsland’s performance, we first examined the performance of different combinations of embeddings and classifiers. Selecting the best-performing combination, we examined the performance of TreasureIsland on the L and M sets, and compared it with a set of methods we curated from the literature. We used Gensim’s Doc2vec package in Python to find the document vectors using both DM and DBOW algorithms. We also used Gensim’s ConcatenatedDoc2Vec package in Python for the concatenated DM + DBOW model. For the baseline model TFIDF, we used Gensim’s Dictionary, TfidfModel packages in Python. First, we trained the embedding models Distributed Memory (DM), Distributed Bag of Words (DBOW), Concatenated DM and DBOW model (DM + DBOW), and Term Frequency–Inverse Document Frequency (TFIDF), on the DNA embedding model dataset. The machine learning dataset, as mentioned before, was formed by using 192 genomes in the training set with a total of 2,864 examples and 20 genomes with 643 examples in the validation set. The machine learning dataset was then vectorized for each of the Doc2vec DNA embedding models DBOW, DM, and DM+DBOW using gradient descent from the Doc2vec inference step. The machine learning dataset vectors were also found for the baseline embedding model TFIDF. The task is formulated as a binary classification task, where the labels are 1,0 representing GEI and non-GEI, respectively. A few machine learning classifiers are used that are most commonly seen associated with document classification tasks, such as SVM, Logistic Regression, and k-Nearest Neighbor. To tune the parameters in the machine learning models, we used 10× cross-validation. The evaluations of the classifiers are done based on their overall accuracy, precision, recall, and *F*_1_-score (harmonic mean of precision and recall). The classification task will also help to evaluate the performance of the different DNA embedding models.

#### 2.3.1 Comparing TreasureIsland with other methods

Here, we compare the predictions from TreasureIsland with other GEI prediction models that have previously shown good results: a tool with high precision based on detecting tRNA fragments (Islander), sequence composition-based tools (IslandPath-DIMOB, Sigi-HMM, AlienHunter), and a hybrid tool (IslandViewer4). The reference dataset used for this task is assembled from 20 genomes (originally from the *M* dataset) separated earlier for the machine learning validation data to prevent any biases from the model. A list of the 20 validation genomes is available in Supplementary Materials. The novel GEIs are calculated by counting the unique GEI predictions made by each predictor that is not predicted by any other predictor.

## 3 Results and Discussion

Figure 5 shows that the DBOW + SVM model has the highest precision, recall, *F*_1_ score, and accuracy. Overall, SVM seems to perform best among all other classifiers in Figure 5. Even though DBOW + SVM performs the best in the classification task, it is interesting that the TFIDF + SVM model also performs well in the classification task, showing that word relevance might indeed be a good factor for DNA embedding. In contrast, the Doc2vec DM embedding model seems to perform poorly in the classification task. We suspect that results from TFIDF that are sometimes comparable with the DBOW embedding may be attributed to the fact that the embedding of DNA includes a lower variance of k-mers, and even fewer unseen *k*-mers when trained with enough data.

**Figure 5:**
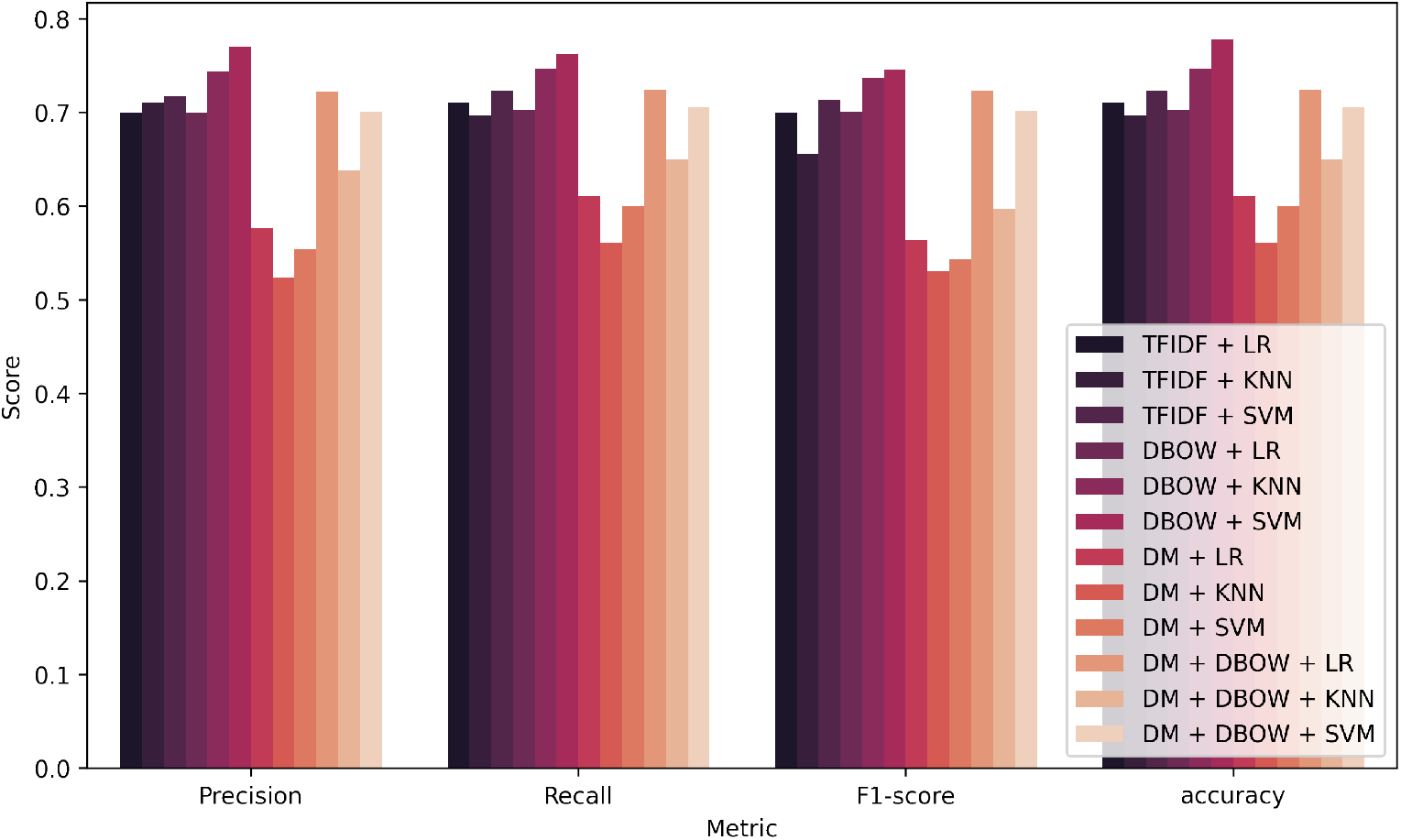
Weighted averaged precision, recall, *F*_1_-score, and accuracy for paragraph vector model Distributed Bag of Words (DBOW) and other baseline representations Term Frequency–Inverse Document Frequency (TF-IDF), Distributed Memory (DM) model and Concatenated DM and DBOW model (DM+DBOW) on classifiers Logistic Regression(LR), Support Vector Machine(SVM), *k*-Nearest Neighbour(KNN). Full results are available in Supplementary Materials.

### 3.1 Evaluation of GEI identification

Although any nucleotide sequence more than or equal to the minimum GEI size *GEI_m_* can be entered to find the GEI regions identified. However, since a typical input would be a whole bacterial chromosome, those are what we used for evaluation. The full list of genomes and results are available in the Supplementary Materials.

We used standard metrics in the field to assess the GEI predictors for the evaluation (Bertelli et al., 2019). The following values were identified based on nucleotide overlaps:

(i) True Positive (TP): The number of nucleotides present in the positive prediction that overlaps with positive reference data. (ii) True Negative(TN): The number of nucleotides outside the positive prediction that overlaps with negative reference data. (iii) False Positive (FP): The number of nucleotides present in the positive prediction that overlaps with negative reference data. (iv) False Negative(TN): The number of nucleotides outside the positive prediction that overlaps with positive reference data.

Based on these values, the following evaluation metrics are used:

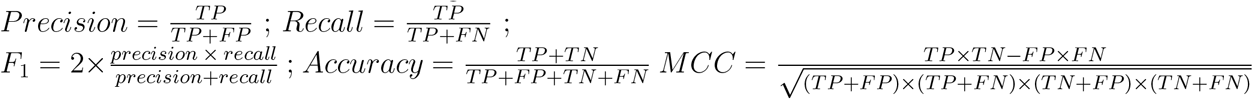

#### 3.1.1 Comparing TreasureIsland with other methods

Table 3 shows that TreasureIsland has the highest recall when compared with other methods. While the precision is lower, the *F*_1_ score, accuracy, and MCC are close to those of Island-Viewer4, and better than the other methods we compared. Figure 6 shows the distribution of MCC and *F*_1_ score across 20 validation genomes in various predictors.

**Table 3:** Performance of TreasureIsland and other baseline GEI predictors on the Benbow validation set. Mean values ± standard error are shown in the table entries.

| Predictor | Precision | Recall | $F_1$-score | Accuracy | MCC | Novel(count) |
| --- | --- | --- | --- | --- | --- | --- |
| TreasureIsland | $0.7863 \pm 0.06$ | **$0.8877 \pm 0.02$** | $0.7890 \pm 0.05$ | $0.8261 \pm 0.05$ | $0.6696 \pm 0.06$ | **531** |
| IslandViewer4 | $0.916 \pm 0.03$ | $0.7474 \pm 0.05$ | **$0.7993 \pm 0.04$** | **$0.8612 \pm 0.03$** | **$0.6836 \pm 0.05$** | N/A |
| IslandPath DIMOB | $0.9108 \pm 0.03$ | $0.4386 \pm 0.06$ | $0.5417 \pm 0.05$ | $0.688 \pm 0.05$ | $0.4373 \pm 0.07$ | 13 |
| SIGI-HMM | $0.9585 \pm 0.02$ | $0.2752 \pm 0.05$ | $0.3878 \pm 0.05$ | $0.6445 \pm 0.05$ | $0.3365 \pm 0.05$ | 3 |
| Islander | **1** | $0.1781 \pm 0.04$ | $0.2603 \pm 0.06$ | $0.6224 \pm 0.06$ | $0.2275 \pm 0.05$ | 0 |
| AlienHunter | $0.6892 \pm 0.05$ | $0.6117 \pm 0.05$ | $0.6017 \pm 0.03$ | $0.6902 \pm 0.04$ | $0.3494 \pm 0.06$ | 346 |

**Figure 6:**
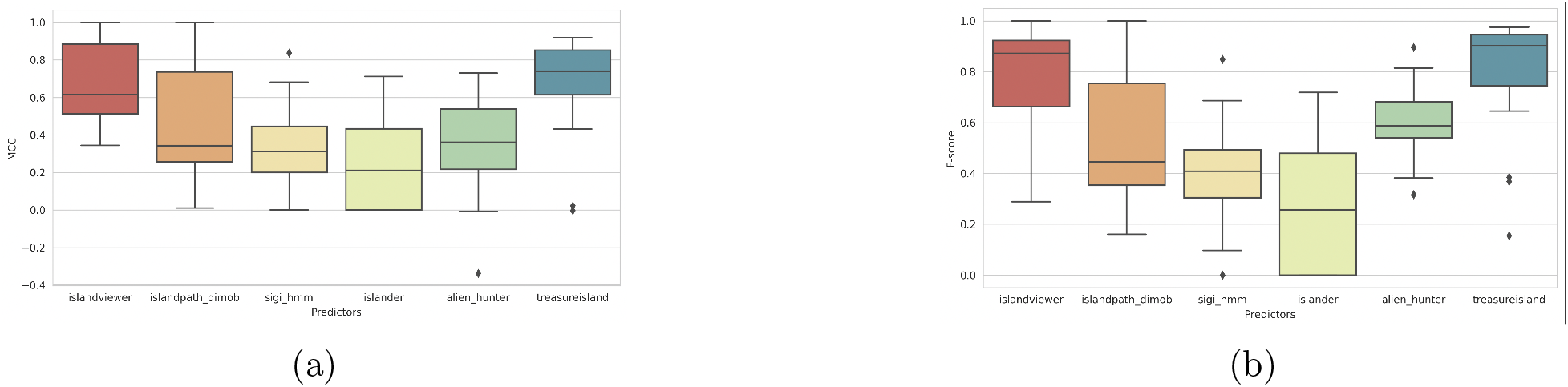
Box plot showing the distribution of performance on 20 genomes from comparative genomics validation data across different predictors.(a) Distribution of MCC values. (b) Distribution of *F*_1_-score values

Additional analysis in Table S16, and Figure S1 shows how changing the *T_u_* can affect the model’s precision and recall, with MCC, accuracy and recall reaching the highest value at 0.80 *T_u_*. The figure shows that at 0.9 *T_u_* TreasureIsland can reach a precision as high as 0.80 while still keeping the recall higher than other models at 0.80, attesting to the strength of recall in our model.

The number of novel predictions made by TreasureIsland is the highest, followed by AlienHunter. IslandViewer4 was excluded from the novel GEI count since it combines the predictions from many of the baseline predictors, leaving no room for novelty. This experiment gives us a good idea about the potential of TreasureIsland to predict GEIs from an input sequence, especially the GEI identification phase in using the models from the first phase. Note that the method performance shown in Tables 3 & 4 are similar to those shown in [8], Box plot showing the distribution of performance on 20 genomes from comparative genomics validation data across different predictors.(a) Distribution of MCC values. (b) Distribution of *F*_1_-score values verifying the veracity of this analysis.

**Table 4:** Performance of TreasureIsland and other baseline GEI predictors on 6 genomes from the L dataset. Mean values ± standard error are shown in the table entries

| Predictor | Precision | Recall | $F_1$-score | Accuracy | MCC |
| --- | --- | --- | --- | --- | --- |
| TreasureIsland | 0.9584 ± 0.02 | **0.9185 ± 0.04** | **0.9341 ± 0.02** | **0.9259 ± 0.03** | **0.8597 ± 0.05** |
| IslandViewer4 | 0.998 ± 0.001 | 0.6691 ± 0.06 | 0.7912 ± 0.05 | 0.8165 ± 0.03 | 0.6837 ± 0.05 |
| IslandPath DIMOB | 0.9976 ± 0.001 | 0.4788 ± 0.06 | 0.6361 ± 0.05 | 0.6998 ± 0.04 | 0.5273 ± 0.05 |
| SIGI-HMM | **1.0** | 0.2048 ± 0.07 | 0.3133 ± 0.09 | 0.5539 ± 0.02 | 0.2716 ± 0.07 |
| Islander | **1.0** | 0.2264 ± 0.06 | 0.3535 ± 0.07 | 0.5600 ± 0.03 | 0.3205 ± 0.04 |
| AlienHunter | 0.7530 ± 0.13 | 0.5703 ± 0.11 | 0.6420 ± 0.12 | 0.7047 ± 0.05 | 0.3979 ± 0.13 |

#### 3.1.2 Experiment on Literature data set

This experiment is done to understand the prediction power of the framework on the literature data set.

##### Experimental setup

The L (Literature) dataset comprises 80 GEIs from 6 genomes. We used the same positive and negative reference GEI data as in [8]. The baseline predictors used in this analysis are IslandViewer 4, IslandPath-DIMOB, Sigi-HMM, Islander and AlienHunter.

##### Results

Table 4, shows the results on the analysis of the curated literature data set. The predictors display a higher precision in this analysis, which means the true positives rate has increased, since the false positive rate is held constant with the same negative data set (from M, the only dataset that has verified true negatives). TreasureIsland shows an improved performance on the literature dataset with highest *F*_1_ score and accuracy, along with recall.

## 4 Conclusions

Here we present TreasureIsland, a document-embedding learning-based framework for predicting genomic islands in unannotated bacterial DNA. TreasureIsland takes unannotated nucleotide sequences as input and uses an unsupervised representation of DNA to classify the GEI and non-GEI regions. We introduce a novel boundary refinement technique to more accurately designate GEI regions. Finally, we provide a new database, Benbow, for training other methods.

We show that TreasureIsland has a high recall and accuracy, and a precision comparable to some of the current baseline predictors. TreasureIsland has the potential to discover novel GEI regions that have not been covered by other predictors. More broadly, we use an unsupervised representation of DNA, which can be helpful in any other DNA-based prediction using machine learning. The advantage of using TreasureIsland is that it does not require any closely related genome sequences nor gene annotations to predict GEIs, which means freshly sequenced unannotated genomes can be used to predict GEI regions.

However, since TreasureIsland does not use any prior information such as gene components in its features, there is a possibility of over-predicting GEIs. These predicted GEIs might fall into the biological grey zone, where we do not know if a region is GEI or not. This problem relates to the broader open-world problem in computational biology, which is the difficulty of obtaining negative training data or to ascertain the veracity of proposed negative training data [24, 25]. While solutions to the problem have been proposed in other ML applications [26, 27], it is still an open problem in many biological applications. Addressing this problem broadly in bioinformatics and biological data science would provide much better classification not only for GEI identification but also in many genomic and biomedical applications where negative data are scarce or unreliable.

## Supporting information

Classification Information

## Acknowledgements

We thank the members of the Friedberg Lab for stimulating discussions.

