## Supplementary materials for "Discovering genomic islands in unannotated bacterial genomes using sequence embedding"

**embedding**

Priyanka Banerjee, Oliver Eulenstein, and Iddo Friedberg

**Table 1:**

Classification results on validation dataset. Weighted averaged precision, recall, f1-score and accuracy for paragraph vector model Distributed Bag of Words (DBOW) and other baseline representations Term Frequency–Inverse Document Frequency (TF-IDF), Distributed Memory (DM) model and Concatenated DM and DBOW model (DM+DBOW) on classifiers Logistic Regression(LR), Support Vector Machine(SVM), K-Nearest Neighbour(KNN).

| Classifier | Precision | Recall | F1-score | Accuracy |
| --- | --- | --- | --- | --- |
| TF-IDF + LR | 0.7 | 0.71 | 0.7 | 0.71 |
| TF-IDF + KNN | 0.7103 | 0.6967 | 0.6559 | 0.6967 |
| TF-IDF + SVM | 0.7174 | 0.7232 | 0.7134 | 0.7232 |
| DBOW + LR | 0.7002 | 0.703 | 0.7013 | 0.703 |
| DBOW + KNN | 0.7437 | 0.7465 | 0.7366 | 0.7465 |
| DBOW + SVM | **0.7801** | **0.7683** | **0.7516** | **0.7683** |
| DM + LR | 0.5759 | 0.6112 | 0.5644 | 0.6112 |
| DM + KNN | 0.5237 | 0.5614 | 0.5303 | 0.5614 |
| DM + SVM | 0.5544 | 0.6003 | 0.5439 | 0.6003 |
| DM + DBOW + LR | 0.7226 | 0.7247 | 0.7234 | 0.7247 |
| DM + DBOW + KNN | 0.638 | 0.6501 | 0.5971 | 0.6501 |
| DM + DBOW + SVM | 0.7005 | 0.7061 | 0.7016 | 0.7061 |

**Table 2:**

Classification results for validation dataset. Mean accuracy with standard error and standard deviation for 10-fold cross validation on different document models Term Frequency–Inverse Document Frequency (TF-IDF), Distributed Bag of Words (DBOW), Distributed Memory (DM) model and Concatenated DM and DBOW model (DM+DBOW) and classifiers Logistic Regression(LR), Support Vector Machine(SVM), K-Nearest Neighbour(KNN).

| Classifier | Mean Accuracy | Standard deviation |
| --- | --- | --- |
| TF-IDF + LR | 0.883726 土 0.00078 | 0.019826 |
| TF-IDF + KNN | 0.818082 土 0.00074 | 0.018863 |
| TF-IDF + SVM | 0.919675 土 0.00069 | 0.017540 |
| DBOW + LR | 0.821590 土 0.00079 | 0.020207 |
| DBOW + KNN | 0.898060 土 0.00071 | 0.018116 |
| DBOW + SVM | **0.922373 土 0.00039** | 0.010042 |
| DM + LR | 0.599285 土 0.00102 | 0.026079 |
| DM + KNN | 0.573330 土 0.00097 | 0.024681 |
| DM + SVM | 0.609153 土 0.00094 | 0.023906 |
| DM + DBOW + LR | 0.811918 土 0.00077 | 0.019744 |
| DM + DBOW + KNN | 0.701823 土 0.00133 | 0.033839 |
| DM + DBOW + SVM | 0.834496 土 0.00094 | 0.023965 |

**Table 3:**

Accuracy of different k-mer sizes on classifiers Logistic Regression(LR), Support Vector Machine(SVM), K-Nearest Neighbour(KNN)

| K-mer size | Window method | | | | | |
| --- | --- | --- | --- | --- | --- | --- |
|  | overlapping window | | | non-overlapping window | | |
|  | LR | SVM | KNN | LR | SVM | KNN |
| 3 | 0.6858 | 0.7698 | 0.6905 | 0.6703 | 0.7045 | 0.6905 |
| 4 | 0.6952 | 0.7514 | 0.6905 | 0.7201 | 0.7543 | 0.6905 |
| 5 | **0.7170** | 0.7618 | 0.7387 | **0.7185** | **0.7589** | **0.7387** |
| 6 | 0.703 | **0.7683** | **0.7465** | 0.6397 | 0.7060 | 0.7231 |
| 7 | 0.6936 | 0.7449 | 0.6983 | 0.6050 | 0.6205 | 0.6983 |
| 8 | 0.6734 | 0.7574 | 0.7014 | 0.6034 | 0.6205 | 0.7014 |
| 9 | 0.6345 | 0.7216 | 0.7092 | 0.5614 | 0.6205 | 0.7092 |

**Table 4:**

Statistical summary of Precision, Recall, Accuracy ,F-score and MCC on 20 comparative genomics validation data for IslandViewer.

|  | Precision | Recall | Accuracy | F-Score | MCC |
| --- | --- | --- | --- | --- | --- |
| mean | 0.916063 土 0.0284 | 0.747487 土0.0504 | 0.861298 土0.0260 | 0.799310 土0.0404 | 0.683650 土 0.0466 |
| Std deviation | 0.127299 | 0.225737 | 0.116667 | 0.180732 | 0.208662 |
| min | 0.513264 | 0.168848 | 0.640599 | 0.288913 | 0.343927 |
| 25% | 0.892089 | 0.592236 | 0.767115 | 0.663117 | 0.514481 |
| 50% | 0.979700 | 0.805478 | 0.887265 | 0.872038 | 0.615965 |
| 75% | 1.000000 | 0.939641 | 0.959056 | 0.923414 | 0.883877 |
| max | 1.000000 | 1.000000 | 1.000000 | 1.000000 | 1.000000 |

**Table 5:**

Statistical summary of Precision, Recall, Accuracy ,F-score and MCC on 20 comparative genomics validation data for IslandPath dimob.

|  | Precision | Recall | Accuracy | F-Score | MCC |
| --- | --- | --- | --- | --- | --- |
| mean | 0.910845 土 0.0346 | 0.438670 土 0.0634 | 0.688004 土 0.0500 | 0.541732 土 0.0563 | 0.437365 土 0.0659 |
| Standard deviation | 0.154936 | 0.283870 | 0.223762 | 0.252121 | 0.295067 |
| min | 0.405393 | 0.087196 | 0.392388 | 0.160405 | 0.010449 |
| 25% | 0.886331 | 0.216054 | 0.481751 | 0.353423 | 0.257158 |
| 50% | 1.000000 | 0.313873 | 0.666233 | 0.445032 | 0.342989 |
| 75% | 1.000000 | 0.606516 | 0.902018 | 0.754770 | 0.736270 |
| max | 1.000000 | 1.000000 | 1.000000 | 1.000000 | 1.000000 |

**Table 6 :**

Statistical summary of Precision, Recall, Accuracy ,F-score and MCC on 20 comparative genomics validation data for SIGI HMM.

|  | Precision | Recall | Accuracy | F-Score | MCC |
| --- | --- | --- | --- | --- | --- |
| mean | 0.958518 土 0.0192 | 0.275289 土 0.0448 | 0.644538 土 0.0504 | 0.387802 土 0.0485 | 0.336574 土 0.0482 |
| Standard deviation | 0.086145 | 0.200502 | 0.225724 | 0.217177 | 0.215953 |
| min | 0.687730 | 0.000000 | 0.345538 | 0.000000 | 0.000000 |
| 25% | 0.982404 | 0.179631 | 0.431621 | 0.304553 | 0.200623 |
| 50% | 1.000000 | 0.257012 | 0.619399 | 0.408925 | 0.312246 |
| 75% | 1.000000 | 0.342234 | 0.857649 | 0.492572 | 0.446129 |
| max | 1.000000 | 0.869269 | 0.977840 | 0.847948 | 0.836298 |

**Table 7 :**

Statistical summary of Precision, Recall, Accuracy ,F-score and MCC on 20 comparative genomics validation data for Islander.

|  | Precision | Recall | Accuracy | F-Score | MCC |
| --- | --- | --- | --- | --- | --- |
| mean | 1.0 | 0.178152 土 0.0437 | 0.622461 土 0.0572 | 0.260320 土 0.0600 | 0.227576 土 0.0543 |
| Standard deviation | 0.0 | 0.195814 | 0.256151 | 0.268356 | 0.243089 |
| min | 1.0 | 0.000000 | 0.052352 | 0.000000 | 0.000000 |
| 25% | 1.0 | 0.000000 | 0.395176 | 0.000000 | 0.000000 |
| 50% | 1.0 | 0.147230 | 0.652176 | 0.256073 | 0.212304 |
| 75% | 1.0 | 0.315781 | 0.817414 | 0.478073 | 0.432833 |
| max | 1.0 | 0.560779 | 0.949407 | 0.718588 | 0.711517 |

**Table 8 :**

Statistical summary of Precision, Recall, Accuracy ,F-score and MCC on 20 comparative genomics validation data for Alien Hunter.

|  | Precision | Recall | Accuracy | F-Score | MCC |
| --- | --- | --- | --- | --- | --- |
| mean | 0.689202 土 0.0465 | 0.611727 土 0.0507 | 0.690282 土 0.0392 | 0.601781 土 0.0332 | 0.349480 土 0.0594 |
| Standard deviation | 0.208311 | 0.227002 | 0.175683 | 0.148804 | 0.266057 |
| min | 0.335959 | 0.224784 | 0.387974 | 0.316796 | -0.337241 |
| 25% | 0.525991 | 0.427686 | 0.532139 | 0.539782 | 0.216711 |
| 50% | 0.719087 | 0.620034 | 0.695500 | 0.586409 | 0.361080 |
| 75% | 0.861424 | 0.789062 | 0.826677 | 0.681537 | 0.539715 |
| max | 1.000000 | 0.948840 | 0.953903 | 0.894233 | 0.729798 |

**Table 9 :**

Statistical summary of Precision, Recall, Accuracy ,F-score and MCC on 20 comparative genomics validation data for TreasureIsland.

|  | Precision | Recall | Accuracy | F-Score | MCC |
| --- | --- | --- | --- | --- | --- |
| mean | 0.786310 土 0.0643 | 0.887720 土 0.0249 | 0.826116 土 0.0532 | 0.789033 土 0.0523 | 0.669550 土 0.0592 |
| Standard deviation | 0.287660 | 0.111403 | 0.238317 | 0.233987 | 0.264908 |
| min | 0.083330 | 0.630812 | 0.088231 | 0.153840 | -0.003635 |
| 25% | 0.692013 | 0.868416 | 0.829950 | 0.744754 | 0.614535 |
| 50% | 0.945386 | 0.914197 | 0.913991 | 0.901541 | 0.740974 |
| 75% | 0.975215 | 0.965298 | 0.947068 | 0.945636 | 0.851513 |
| max | 1.000000 | 1.000000 | 0.964480 | 0.975504 | 0.919817 |

**Table 10 :**

Statistical summary of Precision, Recall, Accuracy ,F-score and MCC on 6 literature validation data for IslandViewer.

|  | Precision | Recall | Accuracy | F-Score | MCC |
| --- | --- | --- | --- | --- | --- |
| mean | 0.998099 土 0.0012 | 0.669120 土 0.0674 | 0.816549 土 0.0319 | 0.791256 土 0.0483 | 0.683736 土 0.0505 |
| Standard deviation | 0.002964 | 0.165295 | 0.078247 | 0.118488 | 0.123808 |
| min | 0.993758 | 0.460823 | 0.745544 | 0.630908 | 0.567502 |
| 25% | 0.996128 | 0.560187 | 0.755955 | 0.716978 | 0.588628 |
| 50% | 1.000000 | 0.650521 | 0.787893 | 0.787236 | 0.641204 |
| 75% | 1.000000 | 0.792481 | 0.876516 | 0.881574 | 0.776972 |
| max | 1.000000 | 0.882978 | 0.926595 | 0.935576 | 0.859175 |

**Table 11 :**

Statistical summary of Precision, Recall, Accuracy ,F-score and MCC on 6 literature validation data for IslandPath Dimob.

|  | Precision | Recall | Accuracy | F-Score | MCC |
| --- | --- | --- | --- | --- | --- |
| mean | 0.997641 土 0.0015 | 0.478883 土 0.0596 | 0.699898 土 0.0414 | 0.636124 土 0.0535 | 0.527399 土 0.0477 |
| Standard deviation | 0.003675 | 0.146042 | 0.101610 | 0.131124 | 0.117001 |
| min | 0.992308 | 0.311434 | 0.528242 | 0.474952 | 0.353072 |
| 25% | 0.995155 | 0.366937 | 0.660759 | 0.535819 | 0.466205 |
| 50% | 1.000000 | 0.482958 | 0.730326 | 0.648781 | 0.549349 |
| 75% | 1.000000 | 0.543012 | 0.748064 | 0.703352 | 0.575676 |
| max | 1.000000 | 0.704652 | 0.818949 | 0.824524 | 0.689684 |

**Table 12 :**

Statistical summary of Precision, Recall, Accuracy ,F-score and MCC on 6 literature validation data for SIGI HMM.

|  | Precision | Recall | Accuracy | F-Score | MCC |
| --- | --- | --- | --- | --- | --- |
| mean | 1.0 | 0.204859 土 0.0690 | 0.553996 土 0.0227 | 0.313323 土 0.0940 | 0.271646 土 0.0675 |
| Standard deviation | 0.0 | 0.169161 | 0.055707 | 0.230444 | 0.165379 |
| min | 1.0 | 0.000000 | 0.489312 | 0.000000 | 0.000000 |
| 25% | 1.0 | 0.092670 | 0.515446 | 0.165735 | 0.200265 |
| 50% | 1.0 | 0.205909 | 0.549358 | 0.340854 | 0.316717 |
| 75% | 1.0 | 0.264649 | 0.579329 | 0.418123 | 0.350516 |
| max | 1.0 | 0.478183 | 0.642486 | 0.646987 | 0.473209 |

**Table 13 :**

Statistical summary of Precision, Recall, Accuracy ,F-score and MCC on 6 literature validation data for Islander.

|  | Precision | Recall | Accuracy | F-Score | MCC |
| --- | --- | --- | --- | --- | --- |
| mean | 1.0 | 0.226429 土 0.0553 | 0.560047 土 0.0330 | 0.353588 土 0.0698 | 0.320506 土 0.0441 |
| Standard deviation | 0.0 | 0.135527 | 0.080844 | 0.171111 | 0.108267 |
| min | 1.0 | 0.066325 | 0.430562 | 0.124400 | 0.194896 |
| 25% | 1.0 | 0.164463 | 0.538586 | 0.282462 | 0.255233 |
| 50% | 1.0 | 0.198437 | 0.564586 | 0.330143 | 0.314516 |
| 75% | 1.0 | 0.257023 | 0.586020 | 0.408652 | 0.346323 |
| max | 1.0 | 0.465685 | 0.677465 | 0.635451 | 0.506708 |

**Table 14 :**

Statistical summary of Precision, Recall, Accuracy ,F-score and MCC on 6 literature validation data for Alien Hunter.

|  | Precision | Recall | Accuracy | F-Score | MCC |
| --- | --- | --- | --- | --- | --- |
| mean | 0.753060 土 0.1291 | 0.570324 土 0.1109 | 0.704716 土 0.0462 | 0.642011 土 0.1219 | 0.397930 土 0.1264 |
| Standard deviation | 0.316267 | 0.271651 | 0.113373 | 0.298699 | 0.309797 |
| min | 0.135759 | 0.022974 | 0.501071 | 0.039298 | -0.176955 |
| 25% | 0.744039 | 0.619367 | 0.681418 | 0.699537 | 0.353387 |
| 50% | 0.880677 | 0.676290 | 0.721528 | 0.758201 | 0.462629 |
| 75% | 0.945899 | 0.706675 | 0.779522 | 0.796902 | 0.612083 |
| max | 0.955114 | 0.729194 | 0.817269 | 0.806581 | 0.660402 |

**Table 15 :**

Statistical summary of Precision, Recall, Accuracy ,F-score and MCC on 6 literature validation data for TreasureIsland.

|  | Precision | Recall | Accuracy | F-Score | MCC |
| --- | --- | --- | --- | --- | --- |
| mean | 0.958398 土 0.0153 | 0.918470 土 0.0444 | 0.925936 土 0.0294 | 0.934105 土 0.0247 | 0.859675 土 0.0509 |
| Standard deviation | 0.037664 | 0.108970 | 0.072233 | 0.060604 | 0.124729 |
| min | 0.885322 | 0.699649 | 0.779616 | 0.813089 | 0.606883 |
| 25% | 0.957428 | 0.939331 | 0.942993 | 0.940805 | 0.890070 |
| 50% | 0.973144 | 0.956577 | 0.955091 | 0.958112 | 0.910251 |
| 75% | 0.976353 | 0.967398 | 0.958987 | 0.964906 | 0.915676 |
| max | 0.989169 | 0.993429 | 0.966575 | 0.973111 | 0.929837 |

**Table 16 :**

Variation of Precision, Recall, MCC, F-score and Accuracy based on changing upper threshold values on 20 comparative genomics validation data for TreasureIsland.

| **Upper Threshold** | **precision** | **recall** | **mcc** | **f-score** | **accuracy** |
| --- | --- | --- | --- | --- | --- |
| 0.5 | 0.7 | 0.93 | 0.61 | 0.75 | 0.79 |
| 0.55 | 0.7 | 0.91 | 0.61 | 0.75 | 0.79 |
| 0.6 | 0.73 | 0.92 | 0.63 | 0.77 | 0.81 |
| 0.65 | 0.75 | 0.9 | 0.64 | 0.78 | 0.81 |
| 0.7 | 0.76 | 0.9 | 0.65 | 0.78 | 0.82 |
| 0.75 | 0.77 | 0.9 | 0.66 | 0.78 | 0.82 |
| 0.8 | 0.78 | 0.88 | 0.67 | 0.79 | 0.83 |
| 0.85 | 0.8 | 0.86 | 0.65 | 0.78 | 0.82 |
| 0.9 | 0.8 | 0.8 | 0.61 | 0.76 | 0.8 |
| 0.95 | 0.84 | 0.72 | 0.59 | 0.73 | 0.79 |

**Figure 1.**

Variation of Precision, Recall, MCC, F-score and Accuracy based on changing upper threshold values on 20 comparative genomics validation data for TreasureIsland. Precision value is noted to increase with higher upper\_thresholds ($T_{u}$), along with decreasing recall value. MCC, f-score and accuracy are seen to reach the highest value at 0.80 $T_{u}$, before declining. Full result is available in Table 16 of Supplementary Materials.

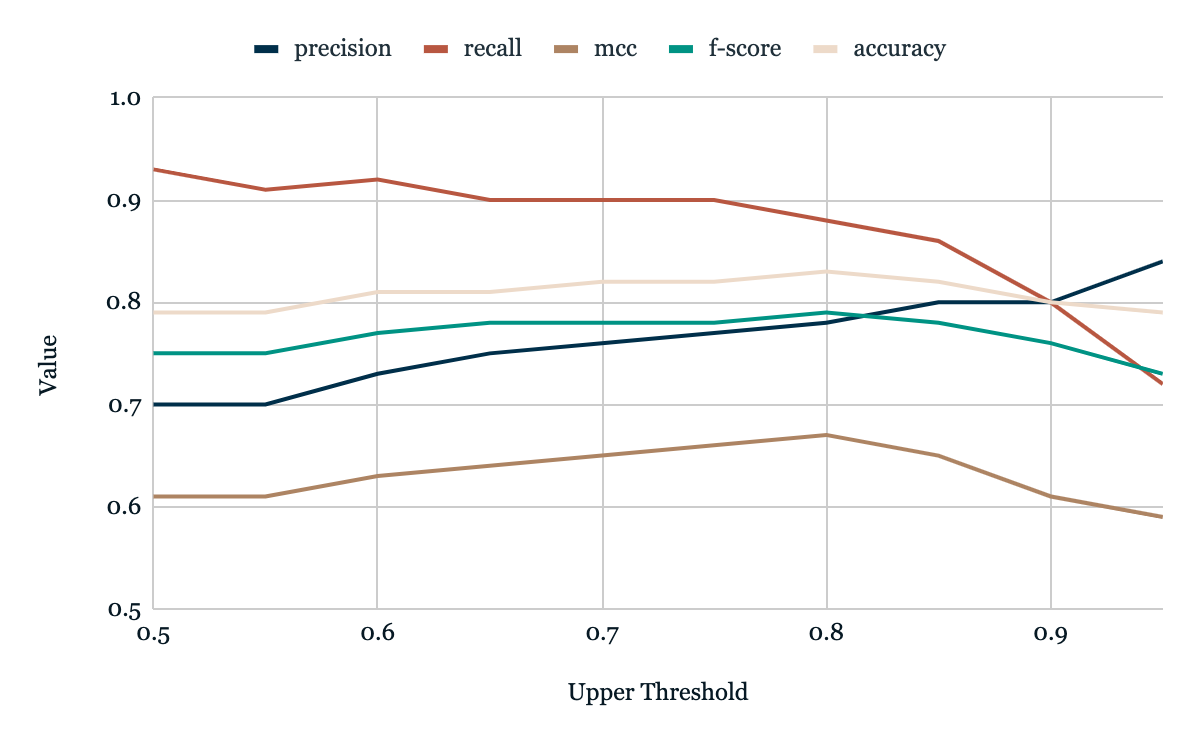
